# On-off transition and ultrafast decay of amino acid luminescence driven by modulation of supramolecular packing

**DOI:** 10.1101/2021.03.23.436384

**Authors:** Zohar A. Arnon, Topaz Kreiser, Boris Yakimov, Noam Brown, Ruth Aizen, Shira Shaham-Niv, Pandeeswar Makam, Muhammad Nawaz Qaisrani, Emiliano Poli, Antonella Ruggiero, Inna Slutsky, Ali Hassanali, Evgeny Shirshin, Davide Levy, Ehud Gazit

## Abstract

It has been experimentally observed that various biomolecules exhibit clear luminescence in the visible upon aggregation, contrary their monomeric state. However, the physical basis for this phenomenon is still elusive. Here, we systematically examine all coded amino acids to provide non-biased insights into this phenomenon. Several amino acids, including non-aromatic, show intense visible luminescence. While lysine crystals display the highest signal, the very chemically similar non-coded ornithine does not, implying a role for molecular packing rather than the chemical characteristics of the molecule. Furthermore, cysteine show luminescence that is indeed crystal-packing-dependent as repeated rearrangements between two crystal structures result in a reversible on-off optical transition. In addition, ultrafast lifetime decay is experimentally validated, corroborating a recently raised hypothesis regarding the governing role of nπ* states in the emission formation. Collectively, our study supports the hypothesis that electronic interactions between molecules that are non-fluorescent and non-absorbing at the monomeric state may result in reversible optically-active states by the formation of supramolecular fluorophores.

## 1. Introduction

In nature, proteins and peptides offer a broad range of characteristics and attributes, which provide an extensive variety of mechanical, electrical and optical properties. These are often derived from the physicochemical properties of the amino acid building blocks that constitute the proteinaceous polymer. In addition to the amino acid content, the arrangement of the protein in its environment, *i.e.*, secondary, tertiary and quaternary structure, can dramatically influence the resulting physical properties. This arrangement is affected by factors such as the solvent, temperature, ionic strength, etc. Recently, several studies described the intrinsic luminescence in the visible range of protein assemblies, while the monomeric protein in solution is not luminescent.^1,2^ Consistently, it is now evident that some amyloids, once assembled, exhibit fluorescence that is not demonstrated in the monomeric state in solution.^3–6^

Monomeric amino acids were subjected to extensive research over the years. Recently, a great emphasis was given to the ability of amino acids to serve as building blocks for the formation of various supramolecular assemblies with attractive features.^7–9^ It was shown that single amino acids in their aggregated form can also produce a fluorescent signal, which is not evident in the monomeric state.^10–13^ Numerous observations suggest that the optical phenomenon of aggregation induced intrinsic fluorescence is broader than originally speculated, as it also applies to many other metabolites and nucleic acids.^14–19^ Circumstantial evidence and argumentation suggested that supramolecular packing is important. It was suggested that the hydrogen bonds and, in specific cases, the aromatic stacking, can underlie these optical properties.^1,20^ Yet, other views have suggested impurities and oxidation as alternative mechanisms, ascribing no significant role to the molecular arrangement.^21^ Indeed, oxidation of aromatic moieties is known to result in the formation of fluorescent products. However, this cannot readily explain the emergence of intrinsic fluorescence upon aggregation of non-aromatic species.^6,19,20^ Overall, the key evidence to distinguish between these options and elucidate the mechanism of intrinsic fluorescence upon aggregation was so far missing. We believe that the reversible transitions between the luminescent and non-luminescent states upon controllable change of aggregate structure is such an evidence. In this work, we aimed at designing and performing such an experiment.

The phenomenon of aggregation-dependent luminescence of proteins is a rapidly evolving field.^1,2,17,22–24,3–5,10–13,16^ Protein aggregates were shown to absorb light at wavelengths above 300 nm and to exhibit a structure-specific fluorescence in the visible range, even in the absence of aromatic amino acids.^6^ A plausible explanation for this phenomenon is the formation of structure-specific supramolecular fluorophores that are permissive to proton transfer across hydrogen bonds.^1,25,26^ Other hypotheses postulate that the fluorescence is associated with electron-hole recombination due to charge transfer between charged amino acids^5,22,27^ or that deep-blue autofluorescence stems from carbonyl double bonds of the protein backbone.^11^ Following these studies, we have decided to look into the intrinsic optical properties of single amino acid and metabolite assemblies. Indeed, we have shown that adenine, phenylalanine, tyrosine and tryptophan exhibit autofluorescence in the visible range upon self-assembly.^10^ This allows the detection of metabolite assemblies within living cells, without any use of external dyes that could interfere with the ability to accurately model a given sample.^28^ Other derivatives of tyrosine were also shown to exhibit luminescence upon aggregation.^29^ Yet, the underlying mechanism of the autofluorescence of these assemblies is controversial, as in all cases it can be attributed to the aromatic moiety. In addition, the explanation of impurities as the source of fluorescence is still a plausible option. Here, we have endeavored to study the optical properties of all 20 coded amino acids in a systematic, non-biased approach. Each amino acid was optically characterized in two states: the original powder form, as obtained from the commercial supplier, and after being dissolved in heated water and allowed to cool down for recrystallization. Aiming to simplify the process of assembly and in order to eliminate as many factors as possible, we avoided altering solvents, salt concentrations, changing pH, etc. The systematic evaluation of all amino acids, before and after recrystallization, provides mechanistic insights into the broad phenomenon of assembly-dependent intrinsic fluorescence.

## 2. Results

First, we examined the optical properties of the crystalline powders of all 20 coded amino acids in the dried powder form. In addition, each powder was dissolved in water at a high concentration (Table S1). In order to completely dissolve the powders, the aqueous solution was heated to 90 °C and vortexed until a clear, transparent solution was achieved. The amino acid solutions were then cooled down gradually to room temperature and incubated for several days to allow the formation of assemblies within the solution. The samples were then lyophilized to attain a dried powder of the amino acid assemblies. The optical properties of these powders were examined as well.

The brightfield and confocal fluorescence images obtained under excitation of 405 nm for all 20 amino acids, as originally received (OS, original Sigma) and after dissolution and reassembly (DR, dissolved and reassembled), as well as quantification of the fluorescence signals, are displayed in **Figure 1** (see **Figure S1** for spectra). For all samples, similar excitation and detection parameters were used. Based on the current measurements there is no systematic trend between the optical properties and the polarity of the amino acids. However, it does appear as though the charged amino acids tend to be characterized by stronger fluorescence whereas those amino acids with hydrocarbon side chains with methyl groups tend to exhibit very weak optical activity.

**Figure 1.**
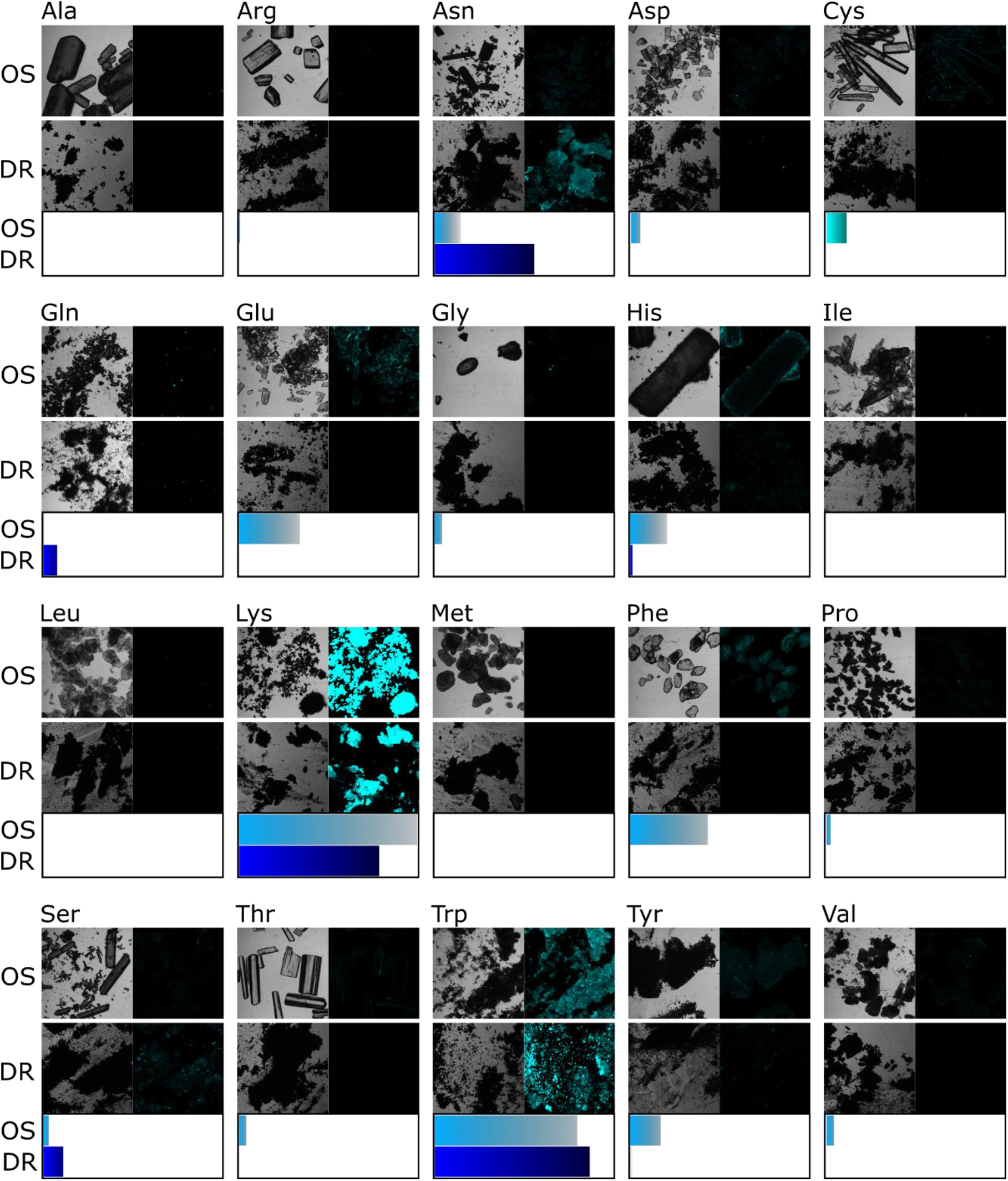
Fluorescence of the 20 coded amino acids. Images of all 20 amino acids, both brightfield (left) and confocal fluorescence at excitation wavelength of 405 nm (right). Amino acids samples as originally obtained from Sigma-Aldrich are displayed in rows marked with OS (Original Sigma). Amino acids dissolved, reassembled and lyophilized are displayed in rows marked with DR (Dissolved and Reassembled). At the bottom of each amino acid panel are the normalized fluorescence signals of the OS sample (light blue) and the DR sample (dark blue).

We then explored the intense fluorescence of L-lysine, which exhibited the brightest signal of all 20 amino acids. We first inquired whether the length of the amine residue chain of lysine, which has four carbons, plays a role in the optical properties of the crystal. For this purpose, we explored the fluorescence of three additional molecules derived from lysine: L-ornithine, L-2,4-diaminobutyric acid and DL-2,3-diaminopropionic acid, which comprise 3, 2 and 1 side chain carbons, respectively **(Figure 2a)**. The rationale of this experiment was to alter the crystal packing by shortening the amine residue, thereby modulating the electronic interactions by varying the distance between molecules, and to examine any effect on the resulting optical properties.

**Figure 2.**
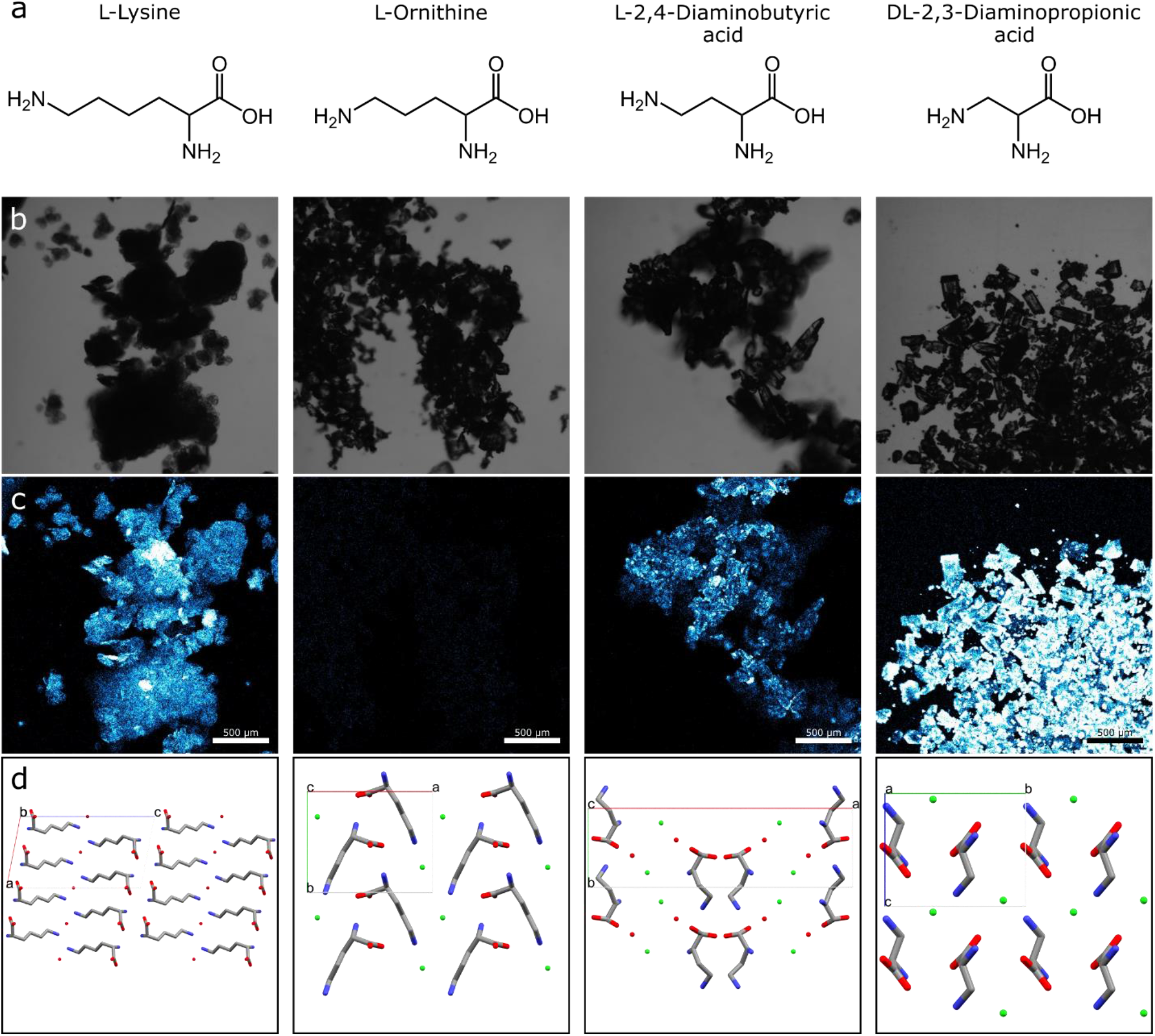
Fluorescence of lysine and its derivatives. **a.** The chemical scheme of lysine and its derivatives comprising shorter carbon chains. **b, c.** Microscopy images showing (b) brightfield and (c) fluorescence (excitation at 405 nm) analyses of the four powders. Pixel color represents the intensity; black–non, blue–dim, white–strong. **d.** Crystalline structures of lysine and its derivatives as determined using PXRD.

All four samples (including lysine) are comprised of small crystalline powders (Figure 2b). The fluorescent signal from each sample was obtained using confocal microscopy, similar to the amino acids shown in Figure 1. Intense fluorescent signals were prominent in the molecules comprising 1-, 2- and 4-carbon chains, while in the 3-carbon chain molecule, ornithine, no visible signal was detected (Figure 2c). The powder X-Ray diffraction (PXRD) patterns of the samples were further examined in order to understand the molecular arrangement of the crystals, which might provide insight into the favorable interactions allowing the crystal fluorescence (Figure 2d). The crystal structures of lysine, ornithine and diaminopropionic acid were previously published.^30–32^ The crystal structure of diaminobutyric acid was determined by the PXRD pattern obtained in this study CCDC Deposition # 1990651 (see Experimental Section). The results revealed the crystal packing of all four samples, allowing molecular inspection of the optical phenomenon. A possible explanation lies in the type of hydrogen bond network that is formed within the supramolecular structure. The NH_2_ group of lysine hemihydrate and diaminobutyric acid, and the NH_3_^+^ of diaminopropionic acid, may interact with a water or chloride ions and donate hydrogen bonds to the molecule complexed in the supramolecular structure. On the other hand, in the case of ornithine, the NH_2_ groups do not form hydrogen bonds that are donated from the amine group to either a chloride ion or a water molecule in the complex. Thus, the differences in the supramolecular packing involving the lysine side-chains, chloride ions and water molecules can lead to differences in the optical properties. We believe that the packing of the crystal is an important factor in determining its optical behavior, much like the properties of microenvironments such as different solvents, ionic strength, polarity or molecular concentration could be significant for structure-function attributes.

Following this line of evidence, we focused on possible interconnections between the optical and structural properties of amino acid powders.

Next, each of the 20 amino acid samples was examined using PXRD to determine the crystalline structure. The results are summarized in Table S2. Most amino acid crystals were composed of the same crystal structure in both the OS and the DR samples. Out of the 20 amino acids, only cysteine and serine displayed different powder diffraction between their respective OS and DR samples, with a single crystal packing observed in each sample. However, both serine samples exhibited some level of fluorescence, while the cysteine DR sample did not show any detectable fluorescence (Figure 1), making the differentiation between the cysteine samples much more straightforward. In addition, the DR serine sample was found to comprise a hemihydrate crystal packing (Table S2), adding another factor of complexity, while both cysteine crystals showed different arrangements of cysteine alone, with four molecules in both unit cells. Hence, we further investigated the OS and DR samples of cysteine **(Figure 3a,b)**, which showed a clear difference in the fluorescent signal (Figure 3c) and exhibited an orthorhombic and monoclinic packing, respectively (Figure 3d,e).^33,34^ Since the process of attaining the OS powder is unknown to us, we have decided to crystallize an orthorhombic cysteine crystal after dissolving the amino acid in order to control the entire procedure. We were able to crystallize both crystal packings, orthorhombic and monoclinic, depending on the crystallization temperature, 4 °C and 25 °C respectively, as the only difference in the crystallization conditions. This allowed a reversible crystallization of both crystal packings, exemplifying the significance of the crystal packing to the optical properties of the crystal. Thus, the fluorescent OS powder (Figure 3f) was dissolved and recrystallized at 25 °C to obtain the non-fluorescent monoclinic DR sample (Figure 3g), and then re-dissolved and recrystallized at 4 °C to form yet again the fluorescent orthorhombic crystals (Figure 3h). To confirm that the fluorescence does not stem from impurities, the non-fluorescent DR sample was washed several times, to remove any impurities that may reside in the supernatant, before recrystallizing as orthorhombic crystals. Although the fluorescent signal of cysteine is relatively low, a difference in the fluorescent signal between the two crystal packings is clearly evident. A comparison of the two structures shows that there is a subtle difference in the proximity of the S-H bonds relative to each other. In the orthorhombic structure, the sulfide groups are packed closer to each other and may facilitate charge transfer, as seen in previous studies.^1,6,22^ It is important to note that the overall macroscopic morphology is dictated by other factors in addition to the packing, as evidenced by different morphologies for an identical packing (compare Figure 3f to 3h), and vice versa (compare Figure 3g to 3h). For this reason, the crystal packing was determined using PXRD for each sample and for every iteration. Morphologically, the recrystallized orthorhombic structures (Figure 3h) are similar to the non-fluorescent monoclinic structures (Figure 3g), and are dissimilar to the relatively large OS crystals (Figure 3f); hence, this affirms that the difference in fluorescence is not derived from any morphological discrepancy. As stated above, the crucial experiment to prove the influence of crystal packing on intrinsic fluorescent formation could be an observation of reversible transitions between the luminescent and non-luminescent states upon controllable change of the crystal structure. We believe that the observed on-off switching of cysteine fluorescence upon changing its crystal packing (Figure 3) represents such an experiment.

**Figure 3.**
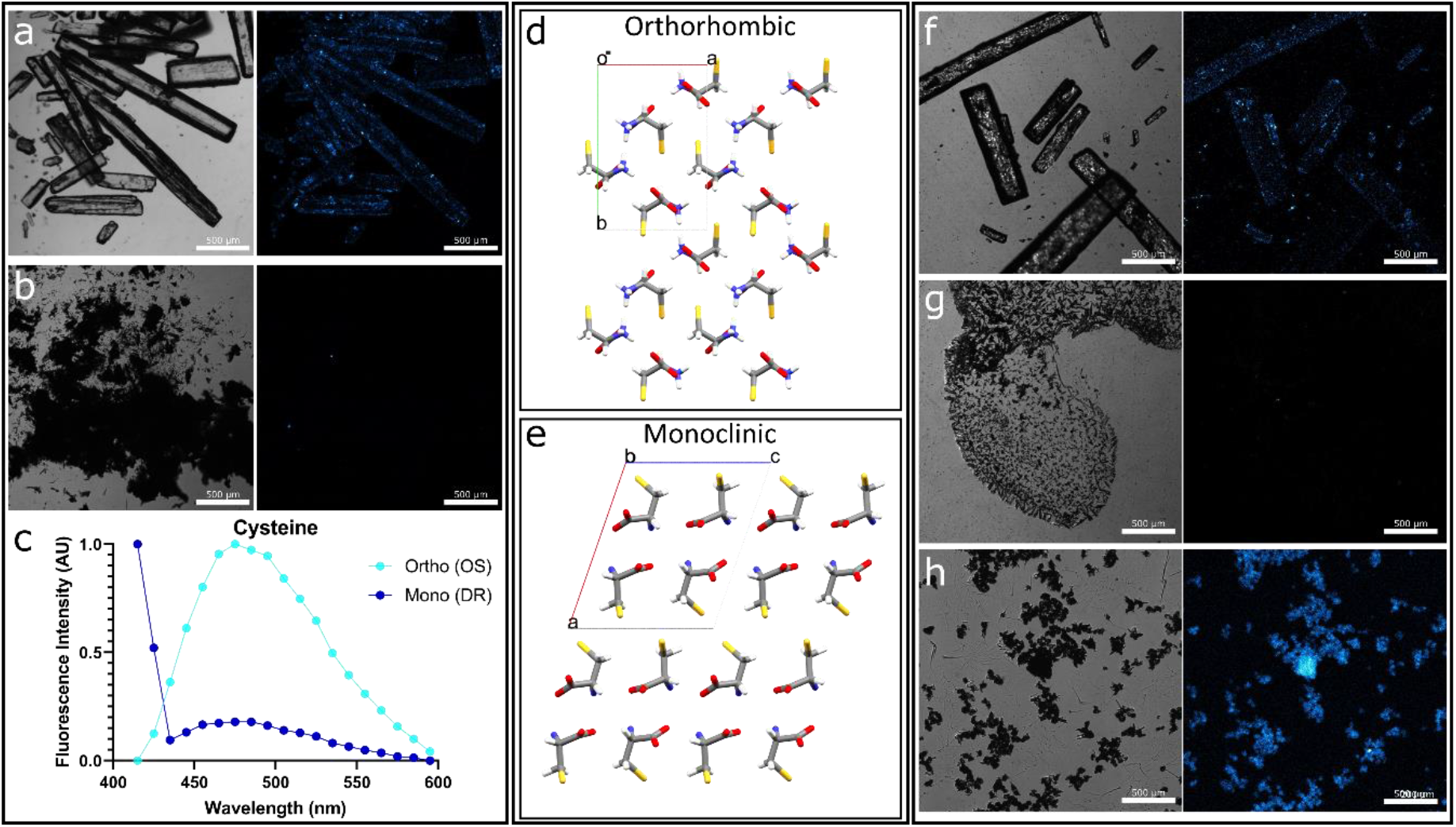
Fluorescence of cysteine. **a, b.** Confocal microscopy images of the cysteine (a) OS sample and (b) DR sample. **c.** The normalized fluorescence intensity of cysteine samples as a function of the emission wavelength (excitation at 405 nm). **d, e.** Crystalline structures of cysteine based on PXRD analysis showing (d) the orthorhombic packing of the OS sample and (e) the monoclinic packing of the DR sample. **f-h.** Confocal imaging of the cysteine (f) OS sample, (g) DR sample and (h) DR sample which was re-dissolved and recrystallized at different conditions to regain an orthorhombic packing. For confocal images, pixel color represents the intensity; black–non, blue–dim, white–strong.

In an attempt to explain the underlying mechanism of the self-assembly induced fluorescence of amino acids, we tried to find a common thread in the aggregation induced fluorescence related literature. First principles quantum chemistry calculations of self-assembly induced fluorescence mainly deal with absorption spectra.^1,19,25^ Without novel absorption bands, no novel fluorescence bands may appear, and the key to the explanation of the presence of new emission properties is the understanding of long wavelength absorption formation. This paradigm was extended in the recent work of Grisanti *et al.*, where, using *ab initio* calculations, fluorescence properties of model peptides aggregates were assessed.^35^ Besides the prediction of the emission in the visible spectral range, a conclusion was made on the time-resolved behavior of the aggregation induced fluorescence. Namely, the presence of an ultrafast fluorescence decay and accompanying spectral diffusion, *i.e.* a gradual fluorescence emission red shift in time, on a sub-picosecond time scale, was described theoretically. This result prompted us to search for experimental indication of an ultrafast component in a self-assembly induced fluorescence system by means of the ultrafast spectroscopy. Taking into account the universal character of this emission, *i.e.* similarity of its spectral properties for a broad range of systems, we chose to work with a known model system – fibrillar structures that are formed as a result of phenylalanine self-assembly. The fluorescence of phenylalanine was previously characterized, and showed the high fluorescent signal, much higher than the unfortunately too weak signal of cysteine.^10^ Thus, by the example of the fibrils made of phenylalanine, we aim at corroborating the presence of an ultrafast fluorescence decay of the self-assembly induced fluorescence.

The self-assembly of phenylalanine in aqueous solution was initiated by cooling a supersaturated solution of phenylalanine (40 mg/ml) at 90 °C to 20 °C, similar to previous work.^10^ This leads to the formation of elongated fibrillary aggregates, as shown by bright-field microscopy **(Figure 4a)**. The phenylalanine fibrils exhibited relative high fluorescence emission in the visible spectral range, which is absent in the monomeric state of phenylalanine, and characterized by the wavelength-dependent Stokes shift, i.e. the spectra measured at longer excitation wavelength exhibited red-shifted emission (Figure 4b). Such a behaviour of the fluorescence emission band is known as the red-edge excitation shift effect and is characteristic for the self-assembly induced fluorescence of amino acids, peptides and proteins.^16,17,27^ The presence of elongated fibrillar structures was detectable by fluorescence lifetime imaging, without the use of any external dyes, only by using the autofluorescence signal of the sample (Figure 4c). The autofluorescence decay curve obtained for the phenylalanine fibrils, revealed the presence of 0.35 and 2.53 ns decay times after fitting to a biexponential model (Figure 4d). However, the temporal resolution, or the width of the instrument response function (IRF) of the fluorescence lifetime imaging setup, which employs the standard time-correlated single photon counting technique, is ~50 ps and does not allow for detecting any ultrafast decay components, which will be completely masked by the 50 ps IRF.^36,37^ Hence, we have utilized the fluorescence up-conversion technique, which provides for sub-picosecond resolution of fluorescence decay (for more information, see ‘Experimental Section’).

**Figure 4.**
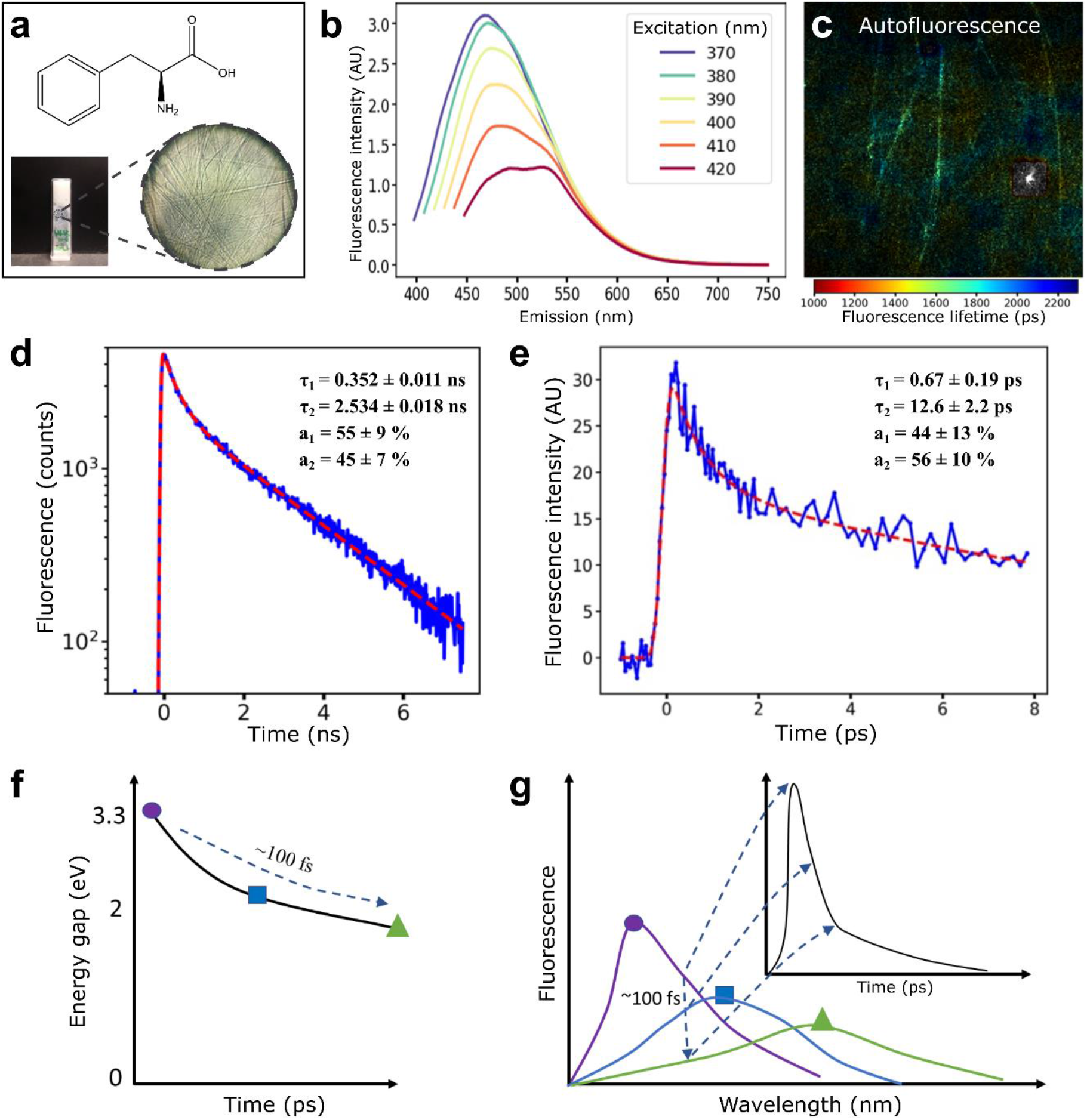
Ultrafast decay. **a.** Chemical scheme and fibrils of phenylalanine (macroscopic and microscopic photos). **b.** Fluorescence emission of phenylalanine assemblies at excitations between 370-420 nm. **c.** Autoluorescence lifetime imaging of phenylalanine fibrils. Excitation was performed in a two-photon regime at 700 nm. **d.** Fluorescence decay curve of the phenylalanine fibrils measured with (sub)nanosecond resolution. **e.** Fluorescence decay curve of phenylalanine fibrils measured using fluorescence up-conversion technique. Excitation and emission were set to 380 and 450 nm, respectively. The inset shows the parameters of fluorescence decay obtained using the biexponential decay model (d,e). **f.** Schematic decrease in the energy gap between the excited nπ* and ground state of the model fibrils. **g.** Schematic of the shift of the aggregation induced fluorescence spectrum with time, demonstrating the presence of the ultrafast decay and spectral migration.

Using fluorescence up-conversion, it was observed that the phenylalanine fibrils exhibited ultrafast decay lifetime component as fast as 0.67 ps. (Figure 4e). The second decay component was characterized by 12.6 ps decay. The presence of the sub-picosecond fluorescence decay is in agreement with the calculation of the recent published work^35^, where the spectral diffusion and relaxation was observed on a few hundred picoseconds time scale.

In the work of Grisanti *et al.*^35^, the ab initio nonadiabatic dynamics simulations were used to reveal characteristic properties of the excited electronic states of model amyloid-like peptides. It was shown that a visible (blue-green) fluorescence could originate from the nπ* states localized on the amide groups. The specific structure of amyloids gives rise to stabilization of some of these states, thus lowering the energy gap between the ground and minimum nπ* state leading to the shift of absorption and fluorescence to the near UV-visible range. Grisanti and co-workers observed that after excitation in the manifold of nπ* states in 10-15 fs all trajectories from different nπ* states reached the lowest excited S_1_ (or nπ*_min_, according to Grisanti *et al.*) state. In the range of 15-40 fs after excitation UV fluorescence from S1 state is observed and is peaked at 3.3 eV (~370 nm). After that, on the time scale of ~100-200 fs, internal vibration redistribution occurs that leads to additional redshift of the emission spectrum, down to 2.0 eV (620 nm) (Figure 4f).

This evolution occurs on the time scale of ~100 fs and is accompanied by the gradual shift of the fluorescence emission from the UV to the blue-green region (Figure 4g). By setting the registration emission wavelength to the blue-green region (in our case, to 500 nm (~2.5 eV)), the ultrafast relaxation along the potential curve shown in Figure 4f can be detected. This process is illustrated in the inset of Figure 4g and corresponds to the fluorescence decay curve presented in Figure 4e. The ultrafast relaxation occurs on the time scale of ~100 fs, corresponding to the non-radiative decay rate of ~10^12^−10^13^ s^−1^, which is, however, lower than in the case of monomeric proteins, where the lack of stabilization of the nπ* states results in the absence of near-UV and visible absorption and blue-green fluorescence.^35^ Moreover, the presence of the ultrafast decay may result in low fluorescence quantum yield of the aggregation-induced emission of proteins (~0.01).^11^ Further relaxation from the lowest energy nπ*_min_ state may then occur on a longer time scale due to the stabilization of the geometry of the aggregate, and this relaxation corresponds to the 1-2 ns fluorescence lifetime, which was observed for the self-assembly induced fluorescence in this paper (Figure 4d) and other previous works.^21^ Hence, by detecting the ultrafast decay of the fibrillar structures formed as the result of phenylalanine self-assembly, the model of Grisanti *et al.* can be extended from peptides to amino acids, therefore elucidating the origin of their enigmatic fluorescence emission upon packing.^35^

## 3. Conclusions

To conclude, in this work we have studied the optical properties of all 20 coded amino acid powders. Each powder was examined as commercially obtained and also after dissolution, reassembly and lyophilization. Surprisingly, lysine exhibited the most intense fluorescence, even though it has no aromatic moieties. Other charged amino acids showed little to no fluorescence. In addition, short-chain derivatives of lysine displayed no correlation between the chain length and their intrinsic fluorescence. This indicates that in some cases, not only the chemical identity of the monomeric molecule dictates the optical properties of the assembly, but also the supramolecular arrangement within the assembly. To substantiate this notion, PXRD analysis together with confocal imaging revealed that the orthorhombic crystal of cysteine is fluorescent, while the monoclinic crystal is not. The recrystallization of the non-fluorescent monoclinic crystals into fluorescent orthorhombic crystals confirms that, in this case, changing the crystal packing is sufficient for conferring optical properties, and that the fluorescence does not stem from contaminations, impurities or oxidation. It is important to note that we do not imply that impurities, oxidation or aromatic interactions are not valid mechanisms that could explain the fluorescence in some cases. However, in addition to those, supramolecular interactions may also affect the optical properties of other biomolecular assemblies, as unambiguously presented here, thus playing a key role in a phenomenon that could be explained by other known mechanisms. In the literature, the phenomenon of self-assembly induced fluorescence is usually addressed for peptides in the context of amyloids optical properties. Several hypotheses have been suggested to explain the mechanism of this effect, and their common motif is that the stabilization of the monomer’s structure by hydrogen bonds in a beta-sheet conformation plays a crucial role in the formation of emitting states. Namely, it was proposed that the peptide aggregation induced fluorescence in the blue-green spectral range is due to (i) delocalization of electrons over a network of hydrogen bonds,^38^ (ii) hydrogen bond-mediated interactions between the amide groups,^39^ (iii) proton transfer across hydrogen bonds,^1^ (iv) decrease in the energy gap between the excited and ground states caused by the influence of the hydrogen bonds on the amide group geometry.^39^ The importance and role of the structure for fluorescence formation in aggregates of non-aromatic peptides has been recently addressed using *ab initio* nonadiabatic dynamics simulations of the excited electronic states.^39^ In addition, previous work has shown that low energy optical excitations and subsequent fluorescence can be induced by charge-transfer excitations.^20,22,40^ Specifically, charge transfer excitations involving sulfur atoms of methionine and the positively charged N-termini of amyloid aggregates were presented.^20^ In the case of cysteine demonstrated in this work, the difference in the molecular packing within the crystal alter the distances between the sulfur atom and the N-termini of the neighboring molecule, which could affect the charge transfer potential. Indeed, these distances are shorter for the orthorhombic crystal in comparison to the monoclinic packing (**Figure S2**). Overall, on the basis of previous research, it can be summarized that the chromophore responsible for the aggregation induced absorption and fluorescence in the visible range is structure-specific, and can be formed either in the absence or presence of aromatic moieties in peptides. Our results demonstrate that even the simplest and most thoroughly investigated molecular systems, such as amino acids, can still serve as the basis for new, intriguing and unknown phenomena. The reversibility of cysteine fluorescence serves as a strong evidence that the molecular arrangement has a crucial role in the observed optical properties.

The prediction of ultrafast decay by Grisanti *et al.* was experimentally confirmed, in an aggregation-induced fluorescence model system of phenylalanine fibrils. Specifically, the presence of the ultrafast decay of the blue-green autofluorescence, are in agreement with the hypothesis regarding the governing role of amyloid structure-stabilized nπ* states in the emission formation. Although the experimental data does not exclude other hypotheses, its relevance to the theoretical calculations can be a step towards understanding the origin of the blue-green emission in amyloids and other systems that recently attracts increased interest, which appear as a result of peptides and amino acid self-assembly. Further understanding of the underlying mechanisms could aid in harnessing the intrinsic properties of supramolecular polymers self-assembled by simple and cost-effective building blocks to develop smart optoelectronic materials.

## 4. Experimental Section

### Amino Acid Reassembly

The original samples from Sigma-Aldrich were dissolved in double distilled water, at a relatively high concentration, depending on the water solubility, as detailed in Table S1. The samples were then heated to 90 °C and vortexed to obtain a clear, transparent solution. The solutions were then incubated at room temperature for a week to allow self-assembly and crystallization. The samples were then lyophilized to remove the water and attain a dry crystalline powder.

### Confocal Imaging

All confocal images were taken using a Leica SP8 Lightning confocal microscope with a Leica Application Suite X (LAS X) software. The samples were excited using a 405 nm laser, with laser intensity set to 50% and the gain set to 500 for all images. The emission range was set between 415 nm to 600 nm. All images were taken at a magnification of 5X.

### Image Analysis

Images were analyzed using Image J. Since we use confocal microscopy, we can assume that the depth (Z) is very small in comparison to the width and height (X and Y). Thus, in these images we can consider the areas rather than the volumes. For each image, we divided the number of “fluorescent” pixels by the number of “black” pixels in the corresponding brightfield image. The threshold for “fluorescent” and “black” pixels were identical for all the images in each experimental set.

### Powder X-Ray Diffraction

X-ray diffraction was collected using a Bruker D8 Discover diffractometer with LYNXEYE EX linear position detector. The diffraction pattern to analyze the amino acid structure was performed in a classical ϑ-ϑ Bragg-Brentano setup. The diffraction patterns were corroborated based on the reported phases in the PDF-4-organics-2019 database. As the crystal structure of L-2,4-diaminobutyric acid was not reported in the literature, a full crystal structure determination was performed (CCDC Deposition # 1990651; Data File S1). The crystalized powder was placed in a 0.7 mm quartz capillary and a full diffraction pattern was collected between 2 and 50 °, step 0.02Å. The capillary setup was employed: Göbels mirrors to obtain a parallel beam, rotating capillary holder. The crystallographic structure determination was performed using the EXPO2014 software.^41^ These EXPO2104 features used were cell indexing (N-TREOR09 algorithm) and the Simulated Annealing Method.

The solution with the lowest cost function was used as model to perform further crystal refinement on the structure using the GSASII software.^42^ The final error indexes were: wR=10.9% and GoF=3.77.

### Cysteine Crystallization

Both cysteine crystal packings were crystallized in double distilled water, at an amino acid concentration of 200 mg/ml. Monoclinic packing was attained as described in the “Amino Acid Reassembly” section above. Orthorhombic packing was attained by incubating the solution over night at 4 °C.

### Phenylalanine fibrils sample preparation

The sample preparation protocol was analogous to the procedures previously used to create self-assembled phenylalanine -aggregates as previously described [Shaham-Niv *et al.*, 2018] using the heat-cool technique. Briefly, L-phenylalanine (Panreac Applichem, CAS 63-91-2, no additional purification) in a concentration of 40 mg/ml was dissolved in distilled water (Millipore-Q) at 90 °C (temperature was controlled by the thermostat Qpod 2e, Quantum Northwest, USA) and was stirred for 1 hour using magnetic stirring for complete dissolving. To obtain self-assembled aggregates, the heated phenylalanine solution was cooled to a room temperature of 23-25 °C under normal conditions in cuvettes or on glass slides, depending on the type of the measurement being performed.

### Steady-state fluorescence measurements of phenylalanine fibrils

Steady-state fluorescence measurements were performed using the FluoroMax-4 spectrofluorometer (HORIBA Jobin Yvon, Japan). Excitation-emission matrices were measured in the 400-750 nm emission range with 1 nm step; the excitation wavelength was varied in the 370-420 nm range with a 10 nm step. The spectral widths of the excitation and emission slits were set to 5 nm. The measurements were carried out in a quartz cuvette with an optical path of 1 cm. The measurements were carried out for samples cooled to room temperature.

### Fluorescence lifetime microscopy (FLIM)

Fluorescence lifetime imaging microscopy (FLIM) with multiphoton excitation was performed using a custom-build multiphoton multimodal microscopy setup. Femtosecond optical parametric oscillator TOPOL-1050-C (Avesta, Russia), providing excitation by signal wave in the 680-1000 nm range, was used as an excitation source. The pulse width of exciting radiation was ~150 fs, with frequency of 80 MHz, average power at the excitation wavelength on the sample was 1 mW. Scanning over the sample was performed using DSC-120 scan head (Becker&Hickl, Germany). Imaging was performed using oil immersion Plan Apochromat 60× objective with NA=1.4 (CFI Plan Apochromat Lambda 60×, Nikon, Japan). Fluorescence decay curves were detected using hybrid GaAsP detector HPM-100-40 (Becker&Hickl, Germany) with sensitivity in 250-720 nm range and instrument response function characteristic time-width of 120 ps. To cut off the excitation, a 680 nm short-pass dielectric filter was used. Both autofluorescence of phenylalanine -fibers and the thioflavin T fluorescence signals were excited at 730 nm.

Fluorescence decay curves were fitted using SPCImage 8.3 software (Becker&Hickl, Germany) after spatial binning (bin size was equal to 5 for ThT-fluorescence measurements and 10 for autofluorescence measurements) by bi-exponential decay law with respect to instrument response function. Average fluorescence lifetime was calculated τ_m_ = (a_1_ τ_1_ + a_2_ τ_2_)/(a_1_ + a_2_), where a_1_, a_2_, τ_1_, τ_2_ are the amplitudes and lifetimes obtained from fit.

### Sub-picosecond fluorescence lifetime measurements

Time-resolved fluorescence emission measurements in the sub-picosecond time range were carried out using a commercially available femtosecond fluorescence spectrometry system FOG100 (CDP Systems, Russia). The samples were excited by 100 fs pulses at 380 nm with a frequency of 80 MHz (second harmonic of Ti:Sapphire oscillator Mai-Tai, Spectra Physics, USA). The fluorescence signal from the sample was focused on a 0.5 mm β-barium borate crystal alongside the fundamental beam (80 fs, 760 nm), acting as a gate pulse for the frequency up-conversion. The gate pulse was delayed by an automatically controlled delay stage. The upconverted light was focused onto the entrance slit of the double monochromator (spectral resolution < 1.5 nm) and was detected by a photomultiplier tube. The reproducibility of the measurement was checked by 10 times, measuring decay trace. Special rotation cuvette unit was used to avoid photodegradation of the sample.

Similar to steady-state fluorescence and microscopy measurements, a phenylalanine sample in a liquid state, heated to a temperature of 90 °C, was poured into a cuvette and was cooled down to room temperature for 15-20 minutes. The measurements were carried out when the sample was turbid. Fluorescence decay curves were fitted by biexponential decay law with respect to instrument response function fitted as Gaussian function. Data analysis on the sub-picosecond decay curves were performed using custom-built Python scripts using LmFit, Matplotlib, Numpy, Pandas libraries.

[CCDC 1990651 contains the supplementary crystallographic data for this paper. These data can be obtained free of charge from The Cambridge Crystallographic Data Centre via www.ccdc.cam.ac.uk/data_request/cif.]

## Supporting Information

Supporting Information is available online or from the author.

## Acknowledgements

Z.A.A. and T.K. contributed equally to this work. We thanks the all our lab members for the fruitful discussions.

## Author contributions

Z.A.A., T.K., S.S.-N., and E.G. conceived and designed the experiments. Z.A.A., T.K., N.B., R.A., S.S.-N., P.M., M.N.Q., E.P., A.R., I.S., A.H., E.S. and D.L. planned and performed the experiments. Z.A.A., T.K., and E.G wrote the manuscript. D.L. performed PXRD experiments and analysis. All authors discussed the results, provided intellectual input and critical feedback and commented on the manuscript.

---

The basis for aggregation-induced fluorescence in the visible range is elusive. Here, we systematically examine the optical properties of all coded amino acids. Several amino acids show intense visible fluorescence. Cysteine exhibit fluorescence that is dependent on the crystal packing, as repeated rearrangements between crystallographic polymorphs resulted in a reversible on-off optical transition. Our study corroborates that electronic interactions between non-absorbing molecules at the monomeric state may result in reversible optically-active states by the formation of supramolecular fluorophores.

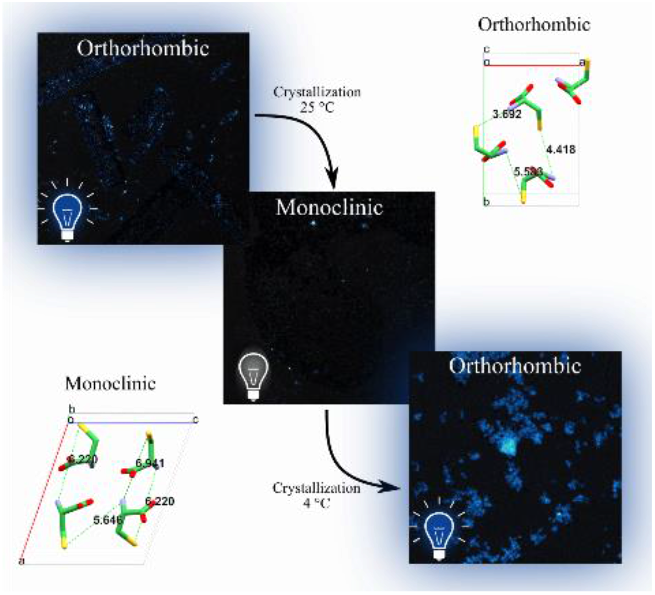

## Supporting Information

**Table S1.** List of amino acids, as purchased from Sigma-Aldrich. DR concentration denotes the concentration used to dissolve and reassemble the amino acid.

| Amino acid | Product name | Product Number | DR concentration [mg/ml] |
| --- | --- | --- | --- |
| Alanine | L-Alanine | A7627 | 200 |
| Arginine | L-Arginine monohydrochloride | 11039 | 1000 |
| Asparagine | L-Asparagine | A-0884 | 200 |
| Aspartic acid | L-Aspartic acid | A9256 | 10 |
| Cysteine | L-Cysteine | 168179 | 200 |
| Glutamine | L-Glutamine | G-3126 | 72 |
| Glutamic acid | L-Glutamic acid | G1251 | 17.2 |
| Glycine | L-Glycine | G7126 | 400 |
| Histidine | L-Histidine monohydrochloride monohydrate | H8125 | 200 |
| Isoleucine | L-Isoleucine | I2752 | 82.4 |
| Leucine | L-Leucine | L8000 | 30 |
| Lysine | L-Lysine | L5501 | 200 |
| Methionine | L-Methionine | M9625 | 100 |
| Phenylalanine | L-Phenylalanine | P2126 | 59.2 |
| Proline | L-Proline | P-0380 | 200 |
| Serine | L-Serine | S4311 | 500 |
| Threonine | L-Threonine | T8625 | 200 |
| Tryptophan | L-Tryptophan | T0254 | 26.8 |
| Tyrosine | L-Tyrosine | T3754 | 2 |
| Valine | L-Valine | V0500 | 100 |

**Table S2.** List of PXRD references (Powder Diffraction File, PDF) for the original sigma powders and the dissolved, reassembled and lyophilized samples. L-Lysine Hemidydrate does not have a PDF number, hence, we include the Cambridge Crystallographic Data Centre (CCDC) number.

| **Amino Acid** | **Sigma Powder (OS)** | **Lyophilized Powder (DR)** |
| --- | --- | --- |
| Alanine | L-Alanine Orthorhombic<br>PDF 02-089-6379<sup>43</sup> | L-Alanine Orthorhombic<br>PDF 02-089-6379 |
| Arginine | L-Arginine Hydrochloride Monoclinic<br>PDF 02-063-2292<sup>44</sup> | L-Arginine Hydrochloride Monoclinic<br>PDF 02-063-2292 |
| Asparagine | L-Asparagine Monohydrate<br>PDF 02-060-1510;<sup>45</sup><br>L-Asparagine<br>PDF 02-095-9836<sup>46</sup> | L-Asparagine Monohydrate<br>PDF 02-060-1510 |
| Aspartic acid | L-Aspartic Acid<br>PDF 02-063-2298<sup>47</sup> | L-Aspartic Acid<br>PDF 02-063-2298 |
| Cysteine | L-Cysteine Orthorhombic<br>PDF 02-073-1763<sup>33</sup> | L-Cysteine Monoclinic<br>PDF 02-063-2319<sup>34</sup> |
| Glutamine | L-Glutamine<br>PDF 02-063-0160<sup>48</sup> | L-Glutamine<br>PDF 02-063-0160 |
| Glutamic acid | L-Glutamic acid<br>PDF 02-063-2343<sup>49</sup> | L-Glutamic acid<br>PDF 02-063-2343 |
| Glycine | L-Glycine Monoclinic<br>PDF 02-063-0180;<sup>50</sup><br>L-Glycine Hexagonal<br>PDF 02-063-0181<sup>51</sup> | L-Glycine Monoclinic<br>PDF 02-063-0180;<br>L-Glycine Hexagonal<br>PDF 02-063-0181 |
| Histidine | L-Histidine Hydrochloride Monoclinic<br>PDF 02-063-0694<sup>52</sup> | L-Histidine Hydrochloride Monoclinic<br>PDF 02-063-0694 |
| Isoleucine | L-Isoleucine Monoclinic<br>PDF 02-063-2412<sup>53</sup> | L-Isoleucine Monoclinic<br>PDF 02-603-2412 |
| Leucine | L-Leucine Monoclinic<br>PDF 02-063-2332<sup>54</sup> | L-Leucine Monoclinic<br>PDF 02-063-2332 |
| Lysine | L-Lysine Hemihydrate<br>CCDC 1488097<sup>30</sup> | L-Lysine Hemihydrate<br>CCDC 1488097 |
| Methionine | L-Methionine Monoclinic<br>PDF 02-063-2419<sup>55</sup> | L-Methionine Monoclinic<br>PDF 02-063-2419 |
| Phenylalanine | L-Phenylalanine<br>PDF 00-063-1180<sup>56</sup> | L-Phenylalanine<br>PDF 00-063-1180 |
| Proline | L-Proline Monohydrate Monoclinic<br>PDF 02-082-9045;<sup>57</sup><br>L-Proline Orthorhombic<br>PDF 02-063-7646<sup>58</sup> | L-Proline Monohydrate Monoclinic<br>PDF 02-082-9045 |
| Serine | L-Serine Orthorhombic<br>PDF 02-063-2442<sup>59</sup> | L-Serine Monohydrate Orthorhombic<br>PDF 02-063-2444<sup>60</sup> |
| Threonine | L-Threonine Orthorhombic<br>PDF 02-063-2447<sup>61</sup> | L-Threonine Orthorhombic<br>PDF 02-063-2447 |
| Tryptophan | L-Tryptophan<br>PDF 00-025-1960<sup>62</sup> | L-Tryptophan<br>PDF 00-025-1960 |
| Tyrosine | L-Tyrosine Orthorhombic<br>PDF 02-063-2456<sup>63</sup> | L-Tyrosine Orthorhombic<br>PDF 02-063-2456 |
| Valine | L-Valine<br>PDF 02-063-2485<sup>64</sup> | L-Valine<br>PDF 02-063-2485 |
| L-ornithine | L-Ornithine hydrochloride<sup>31</sup> | - |
| L-2,4-diaminobutyric acid | L-2,4-Diaminobutyric Acid Hydrochloride<br>CCDC Deposition # 1990651 | - |
| DL-2,3-diaminopropionic acid | (R)-2,3-diammoniopropanoate chloride<sup>32</sup> | - |

**Table S3.** List of Hydrogen bonds lengths (Å) for both the orthorhombic and monoclinic crystal packings.

| Orthorhombic | Monoclinic |
| --- | --- |
| 2.763 (x8), 2.79 (x7), 3.013 (x7) | 2.721 (x3), 2.765 (x2), 2.886 (x4),<br>2.914 (x8), 2.97 (x4) |

**Figure S1.**
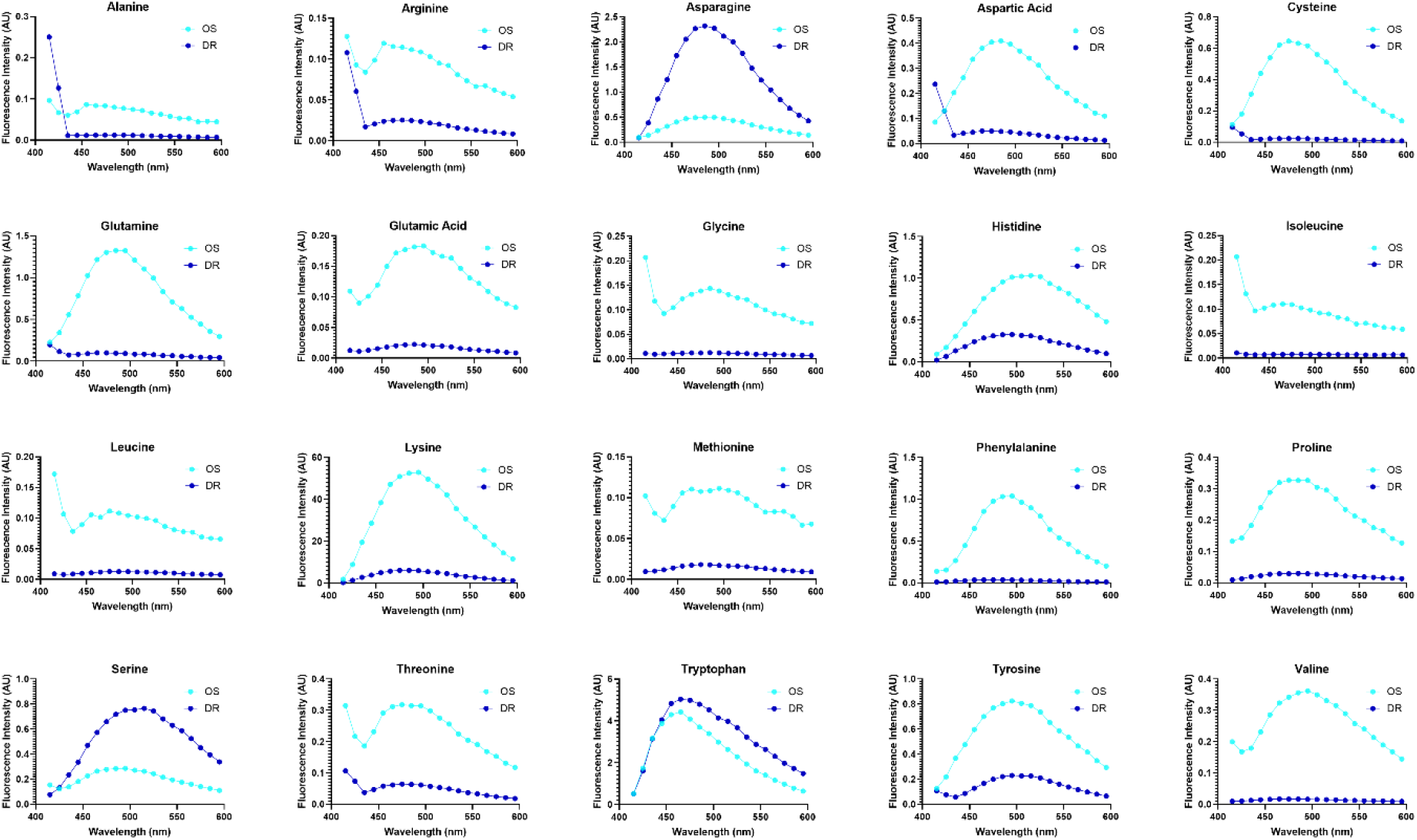
Fluorescence spectra of all 20 amino acids. The fluorescence intensity of all samples as a function of the emission wavelength (excitation at 405 nm), obtained by confocal microscopy.

**Figure S2.**
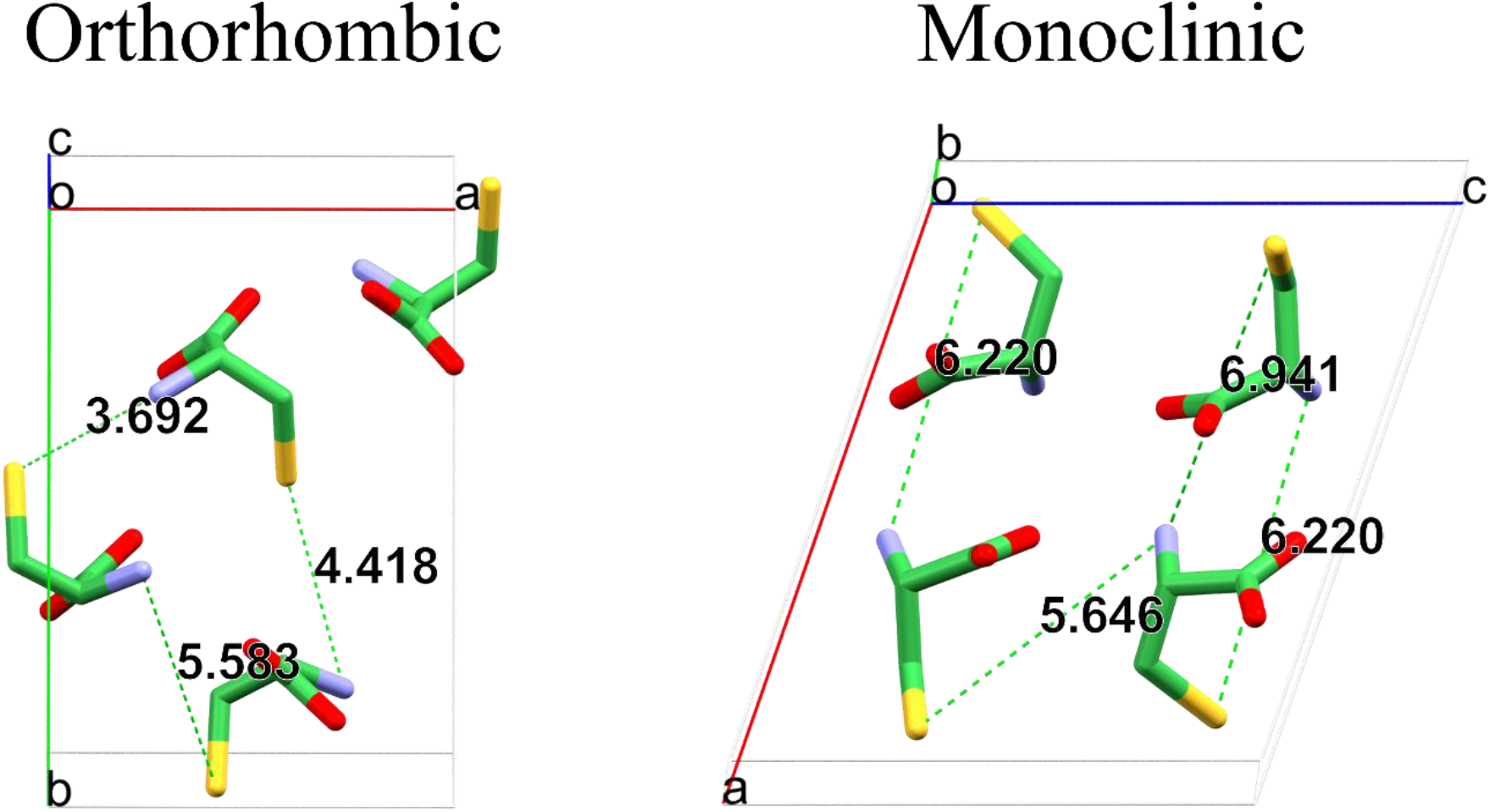
Cysteine crystal structure distances. The measured distances between sulfide atoms and their closest neighboring molecule’s amide. Distances displayed are in Å units.

**Data S1.**

Crystallographic Information File (CIF) for the crystal structure of L-2,4-diaminobutyric acid (CCDC Deposition # 1990651).

